# Dissecting differential signals in high-throughput data from complex tissues

**DOI:** 10.1101/402354

**Authors:** Ziyi Li, Zhijin Wu, Peng Jin, Hao Wu

## Abstract

Samples from clinical practices are often mixtures of different cell types. The high-throughput data obtained from these samples are thus mixed signals. The cell mixture brings complications to data analysis, and will lead to biased results if not properly accounted for. We develop a method to model the high-throughput data from mixed, heterogeneous samples, and to detect differential signals. Our method allows flexible statistical inference for detecting a variety of cell-type specific changes. Extensive simulation studies and analyses of two real datasets demonstrate the favorable performance of our proposed method compared with existing ones serving similar purpose.

## Background

High-throughput technologies have revolutionized the genomics research. The early applications of the technologies were largely on cell lines, for example, by the ENCODE consortium [1]. With the launch of the Precision Medicine Initiative, they have been increasingly applied in larger-scale, population level clinical studies in the hope of identifying diagnostic biomarkers and therapeutic targets. The samples in these studies are often complex and heterogeneous. For example, epigenome-wide association studies (EWAS) profile the DNA methylation in blood samples from a population. In cancer research from large consortium such as The Cancer Genome Atlas (TCGA), genomics and epigenomics signals are measured from solid tumor tissues. The Rush Memory and Aging Project [2] generates a variety of high-throughput data from the postmortem brain samples.

These samples, such as blood, tumor, or brain, are mixtures of many different cell types. The sample mixing complicates data analysis because the experimental data from the high-throughput experiments are weighted average of signals from multiple cell types. In EWAS, the mixing proportions are reported to be confounded with the experimental factor of interest (such as age). The confounding results in many false positive loci if the cell compositions are not properly accounted for [3]. The need to account for sample mixing in the data analysis of complex tissues has gained substantial interests recently, and inspired several methods and softwares, mostly under the context of EWAS studies [3, 4, 5, 6, 7, 8, 9, 10, 11, 12, 13, 14, 15].

One of the most fundamental question in high-throughput data is the differential analysis, e.g., to detect differential expression (DE) or differential methylation (DM) under distinct biological conditions. In complex samples, it is very important to identify cell-type specific changes. Brain tissue, as an example, has a number of distinct cell types, which present highly heterogeneous functions and distinct (epi)genomic profiles [16]. As an illustration, Figure 1(a) shows the gene expression profiles of primary brain cells from rat (data obtained from GSE19380 [17]), where dramatic differences can be observed across cell types. It has also been recognized that distinct cell types get involved in disease pathogenesis and progression with different levels and roles. For example, astrocytes become activated and engaged in neuroinflammatory component, which is related with neurodegeneration process [18, 19, 20]; oxidative damage of microglia, on the other hand, is an important factor for the pathological lesions of Alzheimer’s Disease [21]. Therefore, in analyzing data from complex samples, identifying cell-type specific DE (csDE) and DM (csDM) are important for the understanding of biological or clinical processes, and identifying effective biomarkers for diagnoses and treatment.

**Figure 1:**
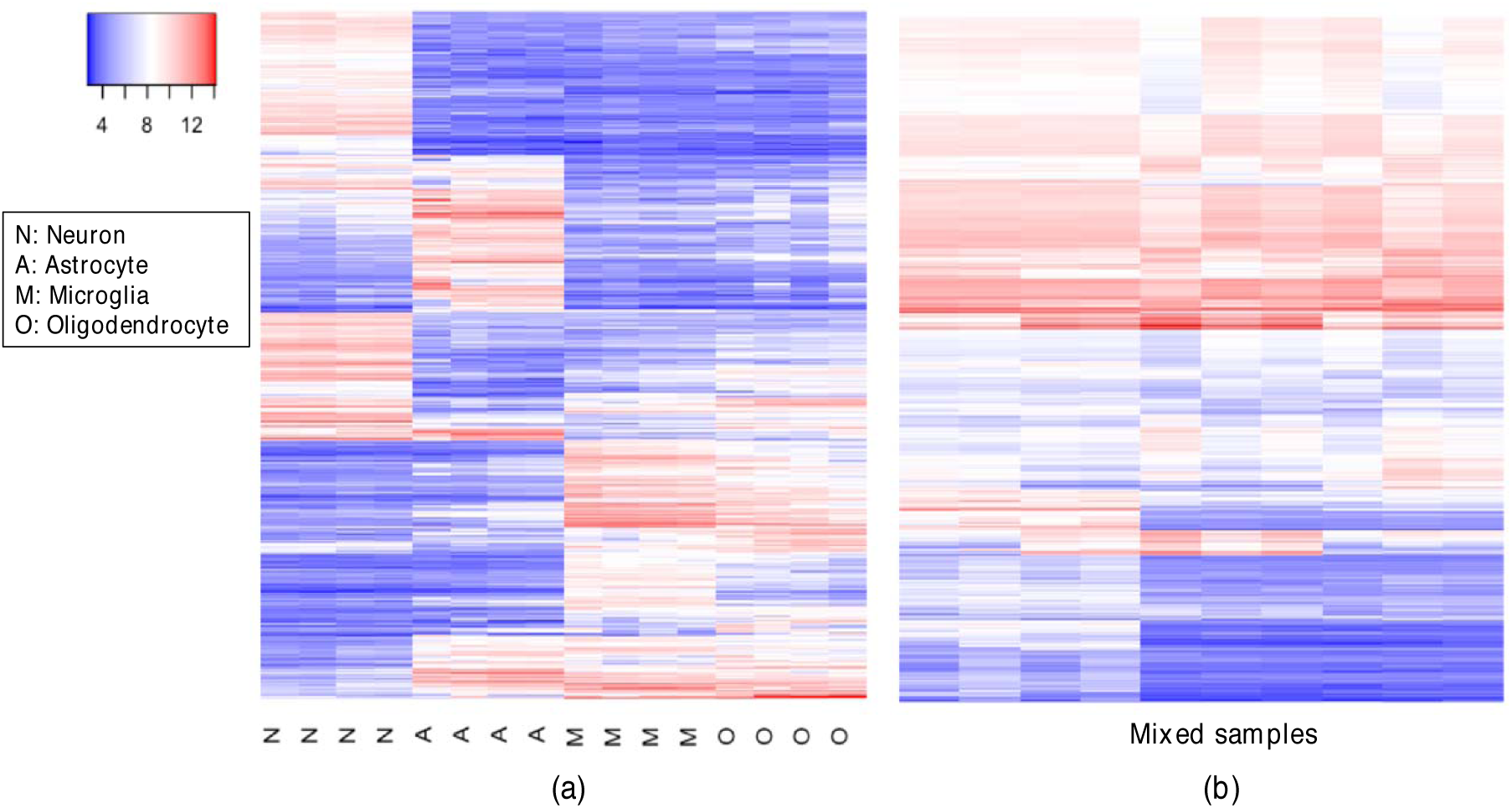
Heatmaps of gene expression profiles for purified rat brain cells and mixed samples. (a) Gene expression from primary brain cell cultures of rat. Purified cell types include neuronal, astrocytic, oligodendrocytic and microglial cultures. (b) Gene expression from RNA mixtures of rat. For both (a) and (b), only the top 1000 most variant genes are presented and the rows have been re-ordered for demonstration purpose. Data downloaded from GSE19380.

There are a number of methods for canonical DE/DM analysis [22, 23, 24, 25, 26]. These methods, however, ignore the cell type mixing, thus directly applying them to the complex sample data will produce undesirable results, including DE/DM due to the change of mixing proportions, or failure to detect DE/DM in specific cell types that is masked in the mixed samples. Figure 1(b) shows the expressions from the mixed samples of primary brain cells in rat. Compared to Figure 1(a), the cell type specific expressions are masked due to cell mixing. For these data, the canonical DE/DM methods will have difficulty to distinguish the cell type specific differences. It is possible to experimentally profile the purified cell types through cell sorting-based technology such as Fluorescent-Activated Cell Sorting (FACS) [27] or Magnetic-Activated Cell Sorting (MACS) [28]. They are, however, laborious and expensive thus cannot be applied to large-scale, population level studies.

Without cell sorting, several statistical methods have been published for identifying cell type specific effects in complex tissue data. These methods usually start with known sample mixture proportions. The *in silico* estimation of mixture proportions is another problem of great interests. Existing methods include reference-based [29, 30, 31, 32, 33, 34], and reference-free methods [4, 17, 35, 36, 37]. Estimating mixture proportions is not the focus of this work, and here we assume the proportions are available, as in the published methods described below. With known mixture proportions, cell-type specific significance analysis of microarrays (csSAM) first estimates the pure tissue profiles by conducting deconvolution on cases and controls separately, and then identifies csDE through permutation tests [33]. The two-step approach (estimating pure profiles and then testing for cell type specific changes) results in lower statistical efficiency and accuracy. Population-specific expression analysis (PSEA) relies heavily on cell-type specific marker genes and use linear models to detect csDE [17]. Other methods including Cell Specific eQTL Analysis [38] and cell-type specific differential methylation detection [5] also use linear model based framework as PSEA does. These methods are designed for specific questions and lack of flexibility to be applied in more general problems.

In this work, we provide a rigorous statistical framework, based on linear model, for characterizing the high-throughput data from mixed samples. Under our model parameterization, the method provides great flexibility for detecting csDE/csDM. A variety of cell type specific inferences can be drawn from testing different linear combinations of the linear model coefficients. Our method, called ***TOAST*** (<u>TO</u>ols for the <u>A</u>nalysis of heterogeneou<u>S</u> <u>T</u>issues), is implemented as an R package and is freely available on GitHub (https://github.com/ziyili20/TOAST). We show (in the Methods section) that all current linear model based csDE/csDM methods are simplified or special cases of TOAST.

## Results and discussion

We conduct extensive simulation and real data analyses to evaluate the performance of the proposed method. We mainly compare the performance of *TOAST* with *csSAM*. In addition, we also include following two procedures for comparison in the simulation studies: *lfc*, which is to directly use the log fold-change of the estimated pure cell type profiles to identify csDE/DM; *absolute diff*, which is to use the absolute difference of the estimated pure cell type profiles to identify csDE/DM. Reference-based method *lsfit* [29] (referred to as “RB”) is used for proportion estimation unless otherwise mentioned. As comparison, reference-free method *deconf* [37] is used under some settings and is referred to as “RF”.

### Simulation

The simulations are focused on evaluating the methods in detecting cell-type specific differential expressions (csDE) from microarray data. We design a series of simulation settings to evaluate the impact of several factors on the accuracy of csDE detection, including signal to noise ratio, sample size, cell mixing proportion magnitudes, and proportion estimation accuracy. The simulation procedures are illustrated in Figure 2, and will be described in detail in the Method section. All simulations are conducted under two-group comparison design. In each setting, simulations are run for 100 times, and the results presented in this section are averages of the 100 simulations. Our criteria to evaluate the methods is the abilities to rank true csDE genes above non-DE genes. We compute the True Discovery Rate (TDR), which is the percentage of true positives among various numbers of top-ranked genes. Method with higher TDR is deemed better.

**Figure 2:**
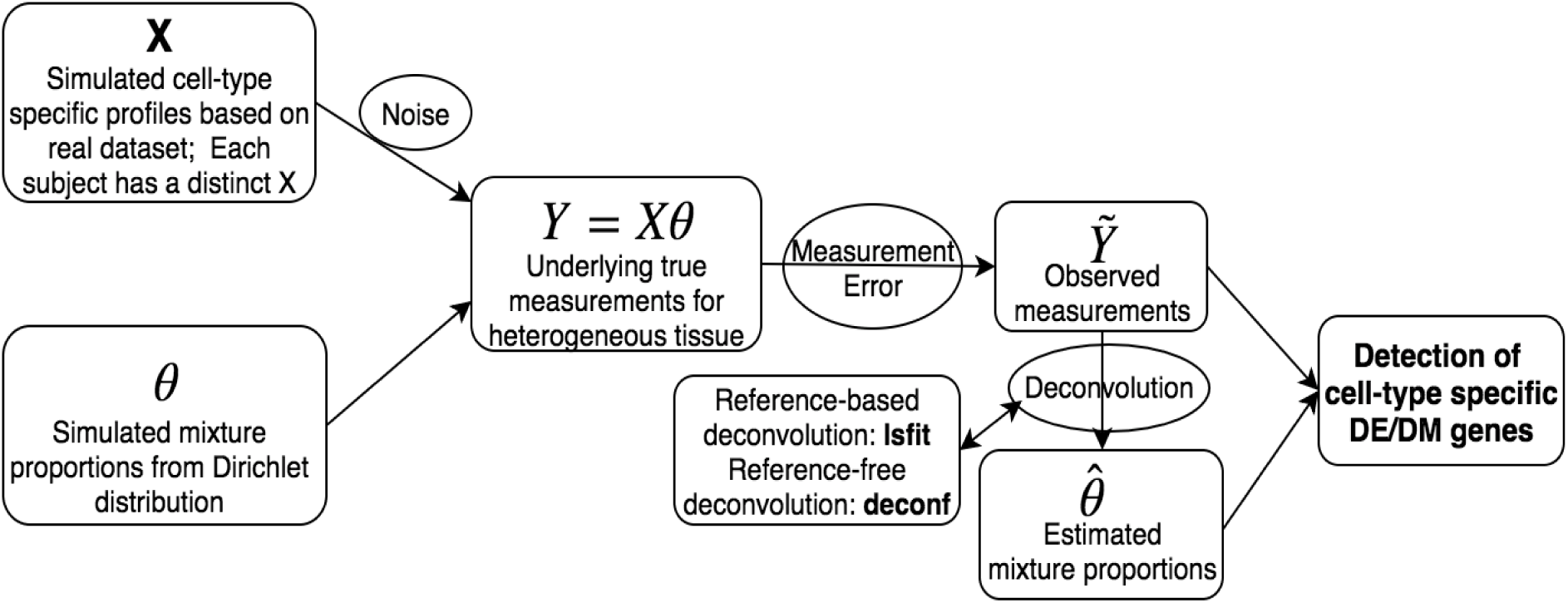
Schematic overview of our simulation study. The general design of our simulation study. From left to right, we first generate simulation datasets, then conduct deconvolution methods to estimate mixture proportions, and lastly apply the proposed method on the synthetic datasets.

We first compare different methods under a typical setting. We assume there are four cell types in the mixture. The data are from two treatment groups with modest sample size (100 samples in case group and 100 samples in control group), and medium noise level. Figure 3 compares the TDR curves for four hypothetical cell types. It shows that using log fold-change of the estimated pure cell type profiles performs very badly and can barely detect true DE genes. Absolute difference is in the middle of log fold-change and csSAM. csSAM demonstrates much better performance, while TOAST provides the best performance in all four cell types. The improvement over csSAM can be substantial. For example, in cell type 4, the true discovery rate from TOAST is close to 100% in top 200 csDE genes, whereas the rate is barely above 70% for csSAM. Overall, the TDR from TOAST is 10% higher than csSAM. Due to the poor performances from log fold-change and absolute difference, we only focus on the comparison of csSAM and TOAST hereafter.

**Figure 3:**
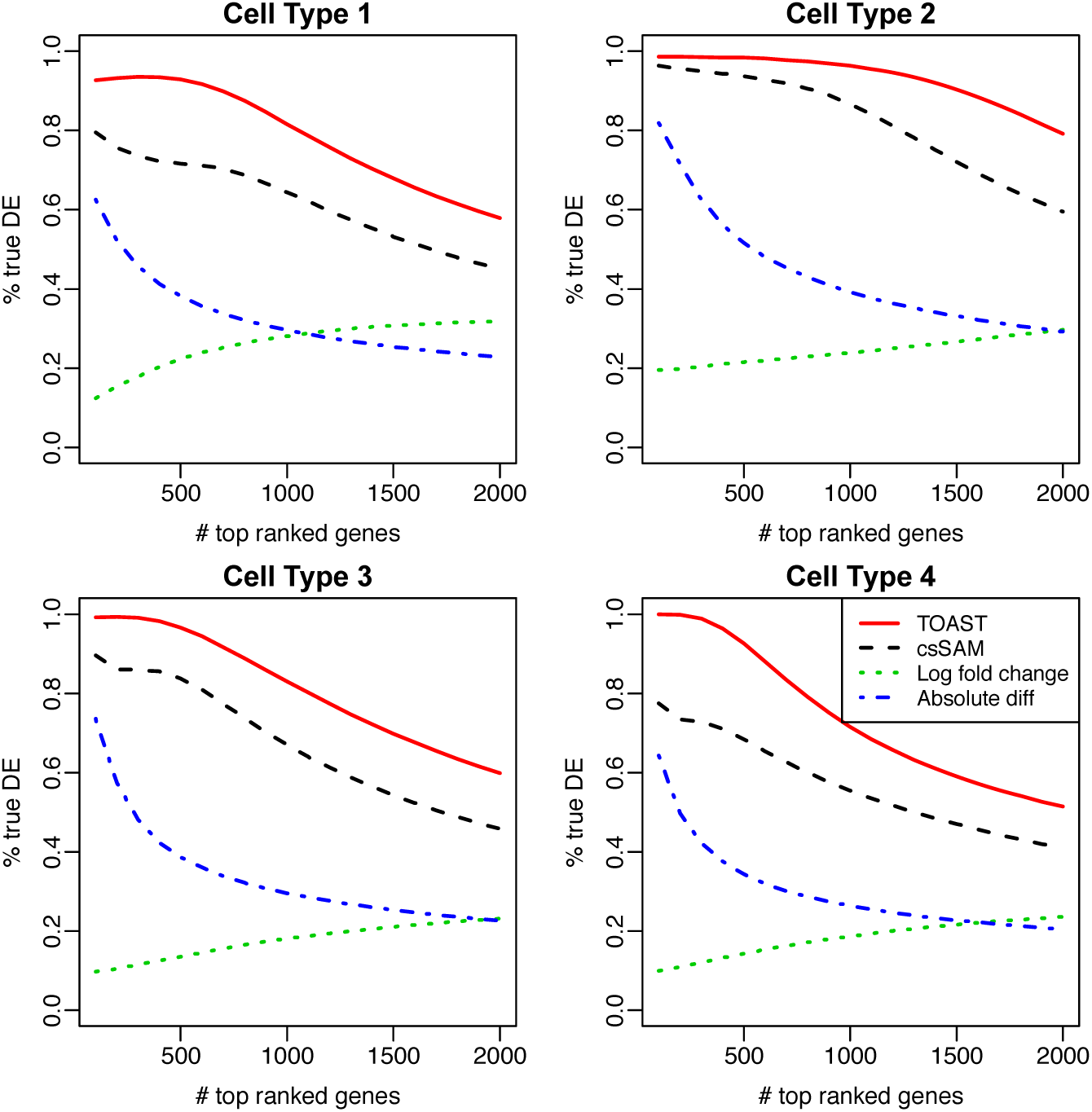
Comparison of csDE detection accuracy from simulation. Shown are the True Discovery Rate (TDR) curves, which plot the proportions of true discovery among top-ranked genes against the number of top-ranked genes. Methods under comparison include TOAST, csSAM, log fold change (lfc), and absolute difference.

#### Impact of noise level and sample size

The noises in the data can come from two sources: (1) biological variation: the variation of pure cell type profiles among different sample; and (2) technical noise: the measurement error. Here we investigate the impact of technical noise level and sample sizes on the performance of the proposed method.

Figure S1 shows the TDR curves from the proposed method under different technical noise levels and sample sizes. We consider noise levels ranging from low (*n*_*sd*_ = 0.1, here *n*_*sd*_ is the parameter controls the magnitude of measurement error and is described in the Method section.) to high (*n*_*sd*_ = 10). Medium level (*n*_*sd*_ = 1) corresponds to the noise level estimated from the Immune Data (described in the Method section), and is close to typical real data observations. We find that technical noise level has substantial influence on the performance of TOAST. When noise level is low, 50 samples in each group is enough for good performance. When noise level is very high, the method suffers significantly especially when sample size is small. In this case, larger sample size (500 samples in each group) can substantially improve the performance.

#### Impact of biological variation

We further evaluate the impact of biological variation (the within-group variation of pure cell type profiles between subjects). Figure S2 shows the TDR curves from the proposed method under different levels of biological variation (controlled by *n*_*ref*_, described in the Method section). *n*_*ref*_ = 1 corresponds to the biological variance level estimated from the the Immune Data. Similar to the technical noise, we find that the biological variation also has substantial impact on the performance of TOAST. This is expected, since greater biological variation reflects higher heterogeneity in samples, which makes it more challenging to detect csDE/csDM. Different from *n*_*sd*_ which multiplies by the variance of a normal distribution, *n*_*ref*_ multiples by the standard deviation of a log-normal distribution. This leads to the results in Figure S2 that, a small change in *n*_*ref*_ can substantially impact the variation of simulated reference panel. When biological variation is small (*n*_*ref*_ = 0.1), the true discovery rate among the 2000 top ranked genes is higher than 0.9 for all four tissues. But when biological variation is large (*n*_*ref*_ = 2), less than 50 percent of the 2000 top ranked genes are true.

#### Impact of proportion magnitude

Results in Figure S1 show that the orders of TDR curves from different hypothetical cell types are similar under all simulation settings, with the red curves being the highest in most cases. Further investigation reveals that this order is related to the abundance of each cell type in the mixture. As described in the Method section, the simulated proportions are based on the proportion estimation of a real dataset (shown in Figure S6). In general, cell type 2 is the most abundant cell type, while cell type 4 is the rarest. So in simulation, cell type 2 has the best ROC curve on most occasions, and cell type 4 has the worst.

To further evaluate the impact of proportion in csDE detection, we conduct another simulation study with all procedures the same as described in the Method section except the proportion generation step. Instead of using real data estimated proportions, we generate proportions from *Dirichlet*(590, 300, 100, 10) for both cases and controls. This ensures the proportions of the first cell type are around 59% for all subjects, second around 30%, etc. Then the obtained TDRs can be linked to the average proportions of the corresponding cell types. In addition, we run simulations using true proportions as input for TOAST, i.e. no proportion estimation procedure involved, thus the impact of proportion estimation accuracy will be excluded. Figure 4 summarizes the distributions of true DE percentages in the top 2000 genes for four cell types, which have different proportions in the mixture. It shows that the magnitude of proportion is closely related with the detection performance. DE genes in cell types with smaller mixture proportions are much harder to be detected. This is expected, because the changes in cell types that make up a smaller proportion of the tissue have smaller contribution to the overall measurement, and thus are more difficult to be identified. To better detect csDE for rarer cell types, the only effective way is to increase the sample size.

**Figure 4:**
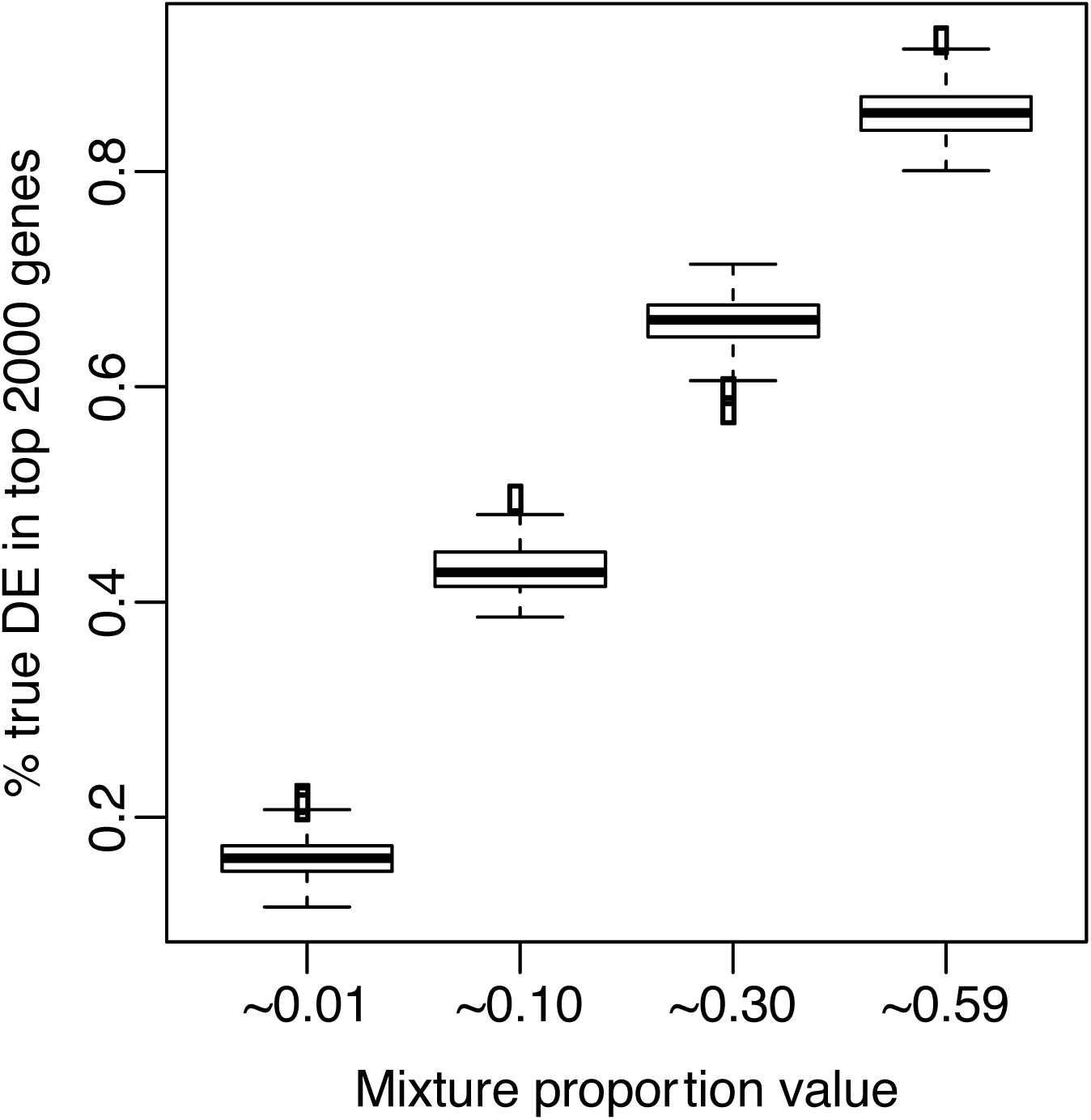
Impact of mixture proportion magnitude on detection accuracy. Boxplot of the detection accuracy (represented by the proportions of true DEs in the 2000 top-ranked genes) by different magnitude of mixture proportion values. ∼ 0.01 means proportion magnitude around 1%.

#### Impact of proportion estimation

Among all the factors, we find the accuracy of proportion estimation has vital impacts on the accuracy of csDE detection. We compared the performances of TOAST and csSAM, using different estimated proportions as inputs. The estimated proportions are from reference-based (RB) and reference-free (RF) methods. We also include the results using true proportions as benchmark.

The TDRs for csDE detection are shown in Figure 5. In all scenarios, TOAST outperforms csSAM. Obviously, using the true proportion gives the best results for TOAST and csSAM. When using the estimated proportions, reference-based estimation provides better results than reference-free estimation. This is as expected, since reference-based method uses extra information (pure cell type profiles), and was reported to produce better proportion estimation [39].

**Figure 5:**
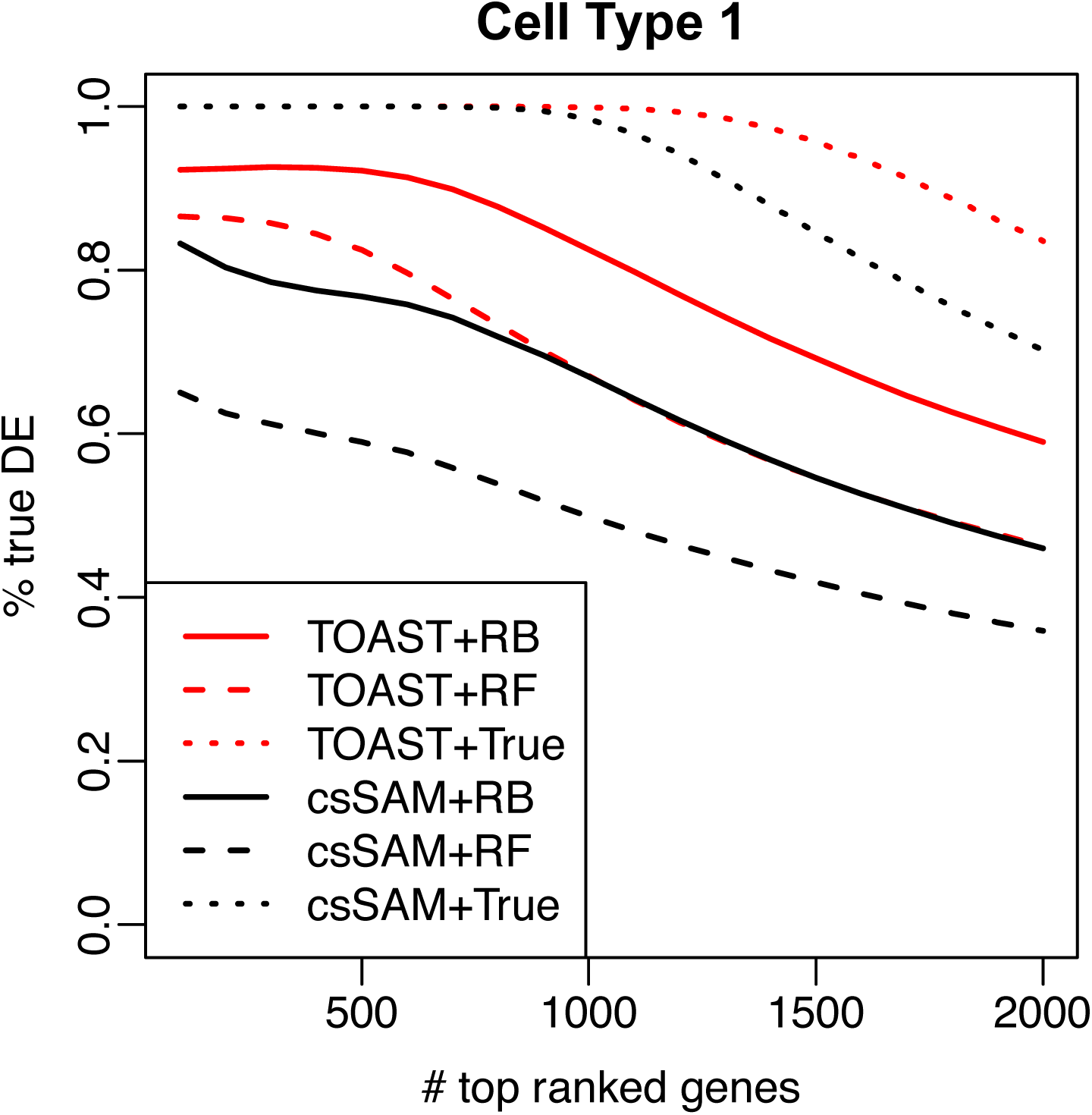
Impact of proportion estimation on TOAST and csSAM. TDR curves comparing TOAST versus csSAM when different deconvolution methods are used as up-stream proportion estimation methods. RB: reference-based. RF: reference-free. True: true proportion.

#### Fitting the model in original or log scale

Historically, gene expression microarray data analyses are primarily performed in logarithm scale. However, under the sample mixing context, the mixing of pure cell type expression values takes place in the raw scale, and the linear relationship will be destroyed if we log-transform the data. This is also the reason why many proportion estimation methods (either reference-free or reference-based) are performed on raw instead of log-transformed data [29, 31, 34, 40].

TOAST fits a linear model for data under the raw scale. We will show that fitting linear model for data in raw scale is statistically justifiable in the Method section. Here, we evaluate the impact of log-transformation on csDE detection by simulation. Figure 6 compares the TDR curves from TOAST for using raw vs. log transformed data in csDE detection for cell type 1, using both reference-based and reference free proportion estimates. It shows that the results from log-transformed the data are reasonable, but using raw scale data gives higher TDR curves with both RB and RF as proportion estimation methods. Results for other cell types lead to the same conclusion. This emphasizes the importance of using raw scale data when performing csDE analysis for mixed data.

**Figure 6:**
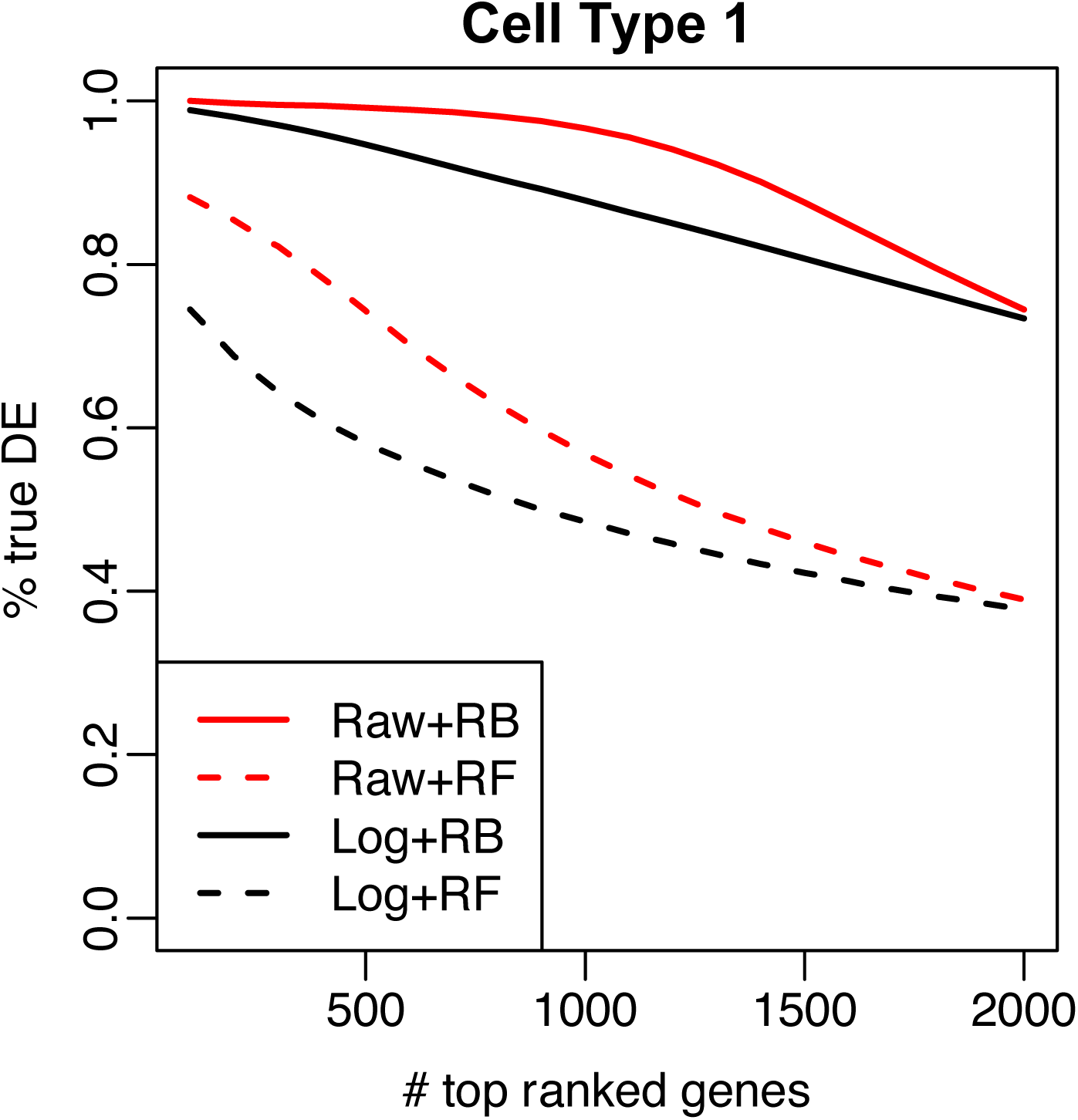
Impact of using raw vs. log-scale data. TDR curvess from TOAST when using raw or log-scale data, and different proportion estimation as inputs for csDE detection. Raw: raw scale data. Log: log scale data. RB: reference-based. RF: reference-free method.

#### Computational performance

TOAST provides superior computational performance since it is based on linear regression. csSAM, on the other hand, relies on permutation procedure and is much more computationally demanding. We benchmark the computational performances of TOAST and csSAM on a laptop computer with 4GB RAM and Intel Core i5 CPU. For a moderate dataset with 20,000 genes and 100 samples in each group, one csDE call from TOAST takes 1.42 seconds, but 378.67 seconds from csSAM. Thus, TOAST is 266 times faster than csSAM.

Overall, the simulation studies demonstrate that our proposed method TOAST provides more accurate and efficient performance in detecting cell type specific changes compared to csSAM. We find that the most important factor in the csDE/csDM detection accuracy is the proportion estimation, which is strongly influenced by the technical noise. Moreover, cell types with lower proportions are more difficult to analyze, since their signal in the mixed data is lower.

## Real data results

### Application to Immune Dataset

In addition to detecting cell-type specific changes among different treatment groups, the proposed method can also detect changes among different cell types in the same group. To demonstrate the functionality of TOAST on this purpose, we obtained a set of gene expression microarray data from NCBI GEO database [29], under accession number GSE11058. The data set includes the gene expression measurements of four immune cell lines (Jurkat, IM-9, Raji, THP-1) and their mixtures. There are four types of mixtures, each with different known mixing proportions. Three replicates are provided for each cell line and mixture. All gene expression data are generated from Affymetrix Human Genome U133 Plus 2.0 Array. This data set is a valuable resource for testing the data analyses method on mixed samples. The differential expression among cell lines can be obtained from pure cell line profiles and used as gold standard to validate the DE calling results from mixed samples. Another advantage is that the known mixture proportions enables us to compare the detection accuracy when using the true versus estimated proportions.

The goal of this data analysis is to detect DE genes for pair-wise comparisons of two different cell lines using the mixture data only, for example, the DE genes between Jurkat and IM-9, IM-9 and THP-1, etc. It is worth noting that we do not compare csSAM in this application because csSAM does not provide the function of detecting DE genes across cell types under the same condition. For each comparison, we define the gold standard based on the pure cell line profiles. We apply *limma* [23] on the pure cell line profiles to call DE genes. The true true DE genes are defined as the ones with (1) the limma p-value smaller than 0.05; and (2) the absolute log fold change greater or equal to 3. The non-DE genes are defined as the ones with limma p-value greater than 0.8. We do not include the genes that have p-values between 0.05 and 0.8 to avoid ambiguity.

Using mixture proportions estimated from reference-based method, we apply TOAST on the mixed data to detect DE genes for all pair-wise comparisons. Figure 7(a) shows the TDR curves for all comparisons. Overall, the proposed method demonstrates good accuracy. The true discovery rate for 3 of the comparisons are over 80% for the top 500 genes. The results for comparing IM-9 and THP-1 is the worst. To further investigate these results, we show the correlation of estimated vs. true proportions in Figure 7(b). The proportion estimation are the worst for IM-9 and THP-1. This partly explains the bad results from IM-9 vs. THP-1 comparison. We further try to use reference-free method to estimate the mixing proportions, and use these estimates as input for DE detection. The results are shown in Figure S3(a). Overall, these results are much worst than using the reference-based proportion estimates. We also use the true proportion as inputs (Figure S4), which yields good accuracy for all comparisons. These results suggest that our proposed method can satisfactorily detect DE genes across cell types from mixed sample data, and that the detection accuracy is highly dependent on the accuracy of proportion estimation. These conclusions are consistent with the simulation results.

**Figure 7:**
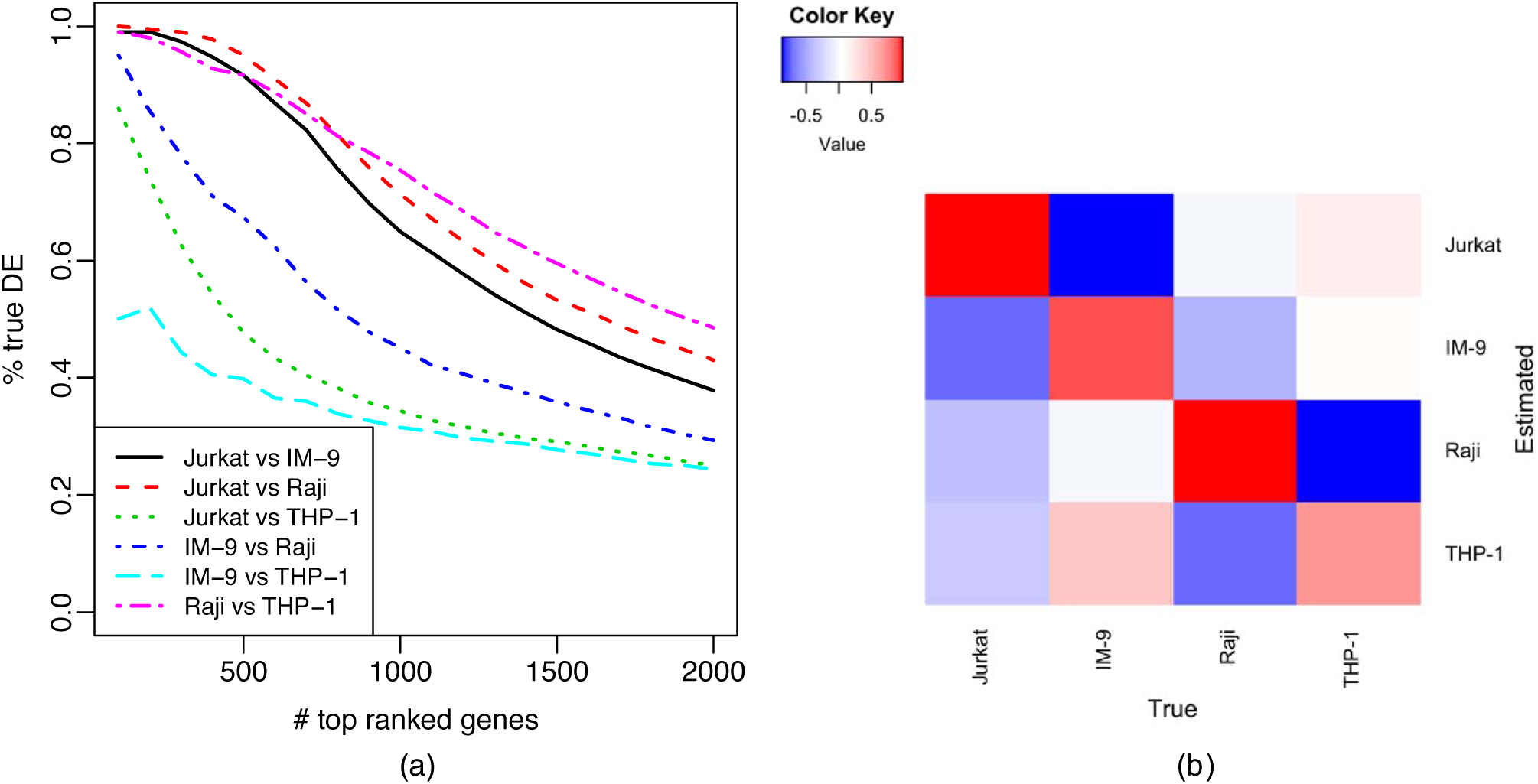
Accuracy of detecting DE across cell types in the Immune Dataset using TOAST. (a) TDR curves for pairwise comparisons of different cell types using TOAST. RB with 1000 reference genes is used to estimate proportions. (b) Heatmap of Pearson’s correlation coefficients between the estimated vs. true mixture proportions.

### Application to Human brain methylation data

To further demonstrate the performance of our method, we analyze a set of human brain DNA methylation data. The data are obtained from GEO database under accession number GSE41826, which include DNA methylation measurements for sorted neuron and glia from post mortem frontal cortex of 29 major depression patients and 29 matched controls. In addition, there are unsorted, whole-tissue measurements from 10 depression cases and 10 matched controls. All data are generated from Illumina Infinium HumanMethylation450 array. In our analysis, we only use the sorted neuron and glia profiles which have matched whole-tissue samples, i.e. 10 cases and 10 controls for sorted pure tissue profiles. The goal of this analysis is to identify differentially methylated CpG (DMC) sites between depression and controls from DNA methylation data of whole tissue samples.

We first construct gold standard DMC using profiles of sorted neuron and glia. We apply Bioconductor package *minfi* [41] to call DMCs between the pure cell type profiles of neuron and glia. True DMC are defined as minfi p-values smaller than 0.05 and the absolute methylation differences greater than 0.05. Non-DM sites are defined as those with minfi p-values greater than 0.8. The DM and non-DM sites are then used as benchmark to evaluate the csDM calling from whole tissue sample data. We estimate the proportions (for neuron and glia) from the whole-tissue DNA methylation data with reference-based deconvolution. The whole-tissue DNA methylation data of 10 cases and 10 controls and the estimated mixture proportions are used as inputs to both TOAST and csSAM. The TDR curves for the results are shown in Figure 8. Overall, these results are not very accurate, because of the complexity of the disease and high inter-individual heterogeneity even for the pure tissue profile. However with the same input data, the proposed method still provides much higher accuracy among the top CpG sites than csSAM. In addition to comparing depression patients versus controls, we also try to detect DMC between other pre-defined conditions. For example, we apply the methods to detect csDM comparing males and females. Figure S5 shows the TDR curves for this comparison. Again, TOAST has higher accuracy of detecting true DMC than csSAM, especially among the top DMs.

**Figure 8:**
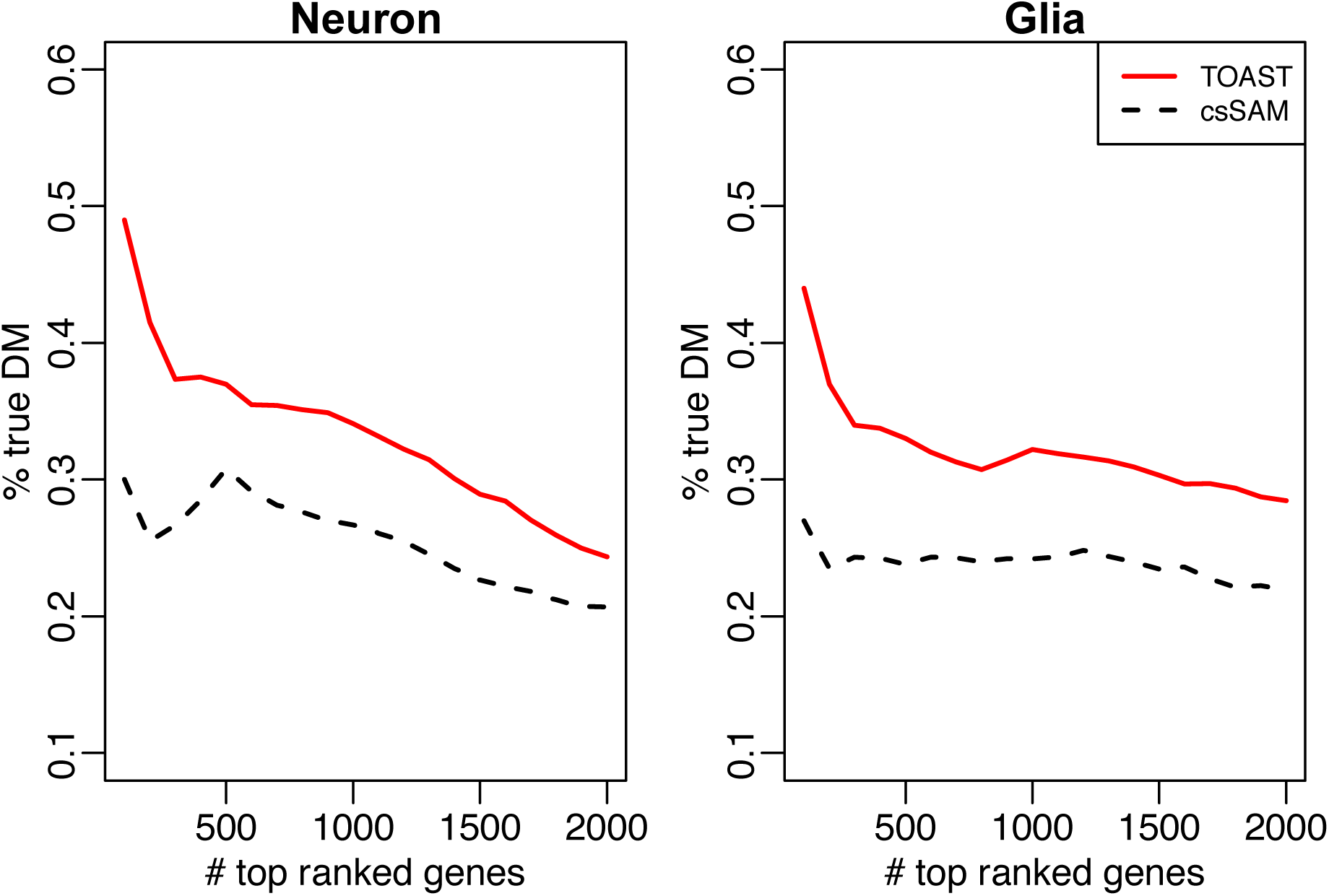
Accuracy of detecting cell-type specific DM in Human brain methylation data. TDR curves for csDM detection from the comparisons of depression patients versus controls in Neuron and Glia, using TOAST and csSAM. RB with 1000 reference genes is used to estimate mixture proportions.

Overall, both real data analyses demonstrate that TOAST can provide accurate and flexible csDE/csDM detection. The results from the human brain data analysis demonstrate that TOAST performs better than csSAM.

## Conclusions

In this work, we develop a general statistical framework to account for sample mixing and detect cell type-specific differential features. The method is based on simple but rigorous statistical modeling, and provides flexible functionalities for testing a variety of cell type-specific differences. A number of previous methods also utilize linear model and are similar in spirit to our method. As shown in the Method section, these methods are simplified version or special cases of our framework. Compared with csSAM, which performs statistical test on deconvoluted pure cell type signals, TOAST can be considered as jointly performing signal deconvolution and hypothesis testing. This results in improved performance, evidenced by simulation and real data analyses. The proposed method can potentially be applied to a number of high-throughput experiments, including but not limited to gene expression microarray data, DNA methylation 450K data, proteomics data, etc.

Based on our simulation studies, the accuracy of mixture proportion estimation is one of the most important factor for the accuracy of csDE/csDM detection. Both simulation and real data analyses show that reference-based estimation performs better than reference-free estimation, which agrees with similar discussions in previous publications [39, 42, 43]. Another advantage of using reference-based deconvolution is that, the corresponding cell types are naturally obtained from reference panel, while it may require an additional step for identifying cell types if reference-free deconvolution is used. Thus, we recommend reference-based method for proportion estimation if possible. Of course, this requires the availability of pure cell type profiles, which could be difficult to obtain for some data. With the accumulation of data, especially from single-cell studies such as the Human Cell Atlas [44], we believe more comprehensive pure cell type profiles will be available as reference in the near future.

Our simulation studies also show that cell types with lower proportions are more difficult to analyze. For those cell types, the only solution to improve detection accuracy is to increase the sample size. This is also related to the problem of choosing *K* (the number of component in the mixture), which is a very difficult model selection question. Complex tissues often contain many (tens even hundreds) cell types if one considers all subtypes of cells. For example, neuron cells can be further divided into many different subtypes such as purkinje cell, granule cell, motor neuron, pyramidal cell, etc. It is impossible to enumerate all cell subtypes in any signal deconvolution or cell type specific differential analysis. In practice, we recommend to put cell types into a few major groups for data analysis. The selection of *K* should be based on biological knowledge.

We anticipate several natural extensions of TOAST. First, the simulation and real data analyses presented in this work are focused on (gene expression or methylation) microarray data, where the data can be approximated by log-normal distribution. The essence of the data modeling and statistical inference from our proposed method can be applied to other types of sequencing data such as RNA-seq, ChIP-seq, or bisulfite-sequencing, even though the detailed model fitting and statistical testing strategies will be different. Secondly, we currently model the effect of covariate in a linear system. It is possible that some covariates (such as age) has a non-linear effect. In this case, we can replace covariates ***Z*** by *f* (***Z***) to model the non-linear effect, where *f* can be a parametric or non-parametric function.

## Method and material

### Data model

Assume data generated from the high-throughput experiments contain measurements for *G* features (genes, CpG sites, etc.) and *N* samples. Denote the measurement for the *g*^*th*^ feature and *i*^*th*^ sample by *Y*_*gi*_. The proposed method is based on the assumption that we have obtained the mixing proportions. The mixing proportions can be experimentally measured [27, 28], or computationally estimated by a number of existing methods [3, 4, 17, 29, 30, 31, 32, 33, 35, 36, 37]. Assume there are *K* “pure” cell types in the mixture, and we have obtained the mixing proportions ***θ*_*i*_** = (*θ*_*i*1_, *θ*_*i*2_,···, *θ*_*iK*_) for sample *i*, (with constraint Σ*k θ*_*ik*_ = 1), our method can perform a variety of differential analysis. Below we use differential expression (DE) as example to illustrate the ideas, though “expression” can be replaced by other measurements such as DNA methylation and the same logic follows.

For the *g*^*th*^ gene in the *i*^*th*^ sample, denote the underlying, unobserved expression in the *k*^*th*^ cell type as *X*_*gik*_. For simplicity of notation, we will drop the subscript *g* in following derivation. Differential expression will be performed one gene at a time (loop over *g*) in the same manner. Let ***Z*_*i*_** be a vector for subject-specific covariates. In a simple two-group comparison without other covariates, ***Z*_*i*_** reduces to a scalar indicator of the non-reference condition (*Z*_*i*_ = 0 for reference, and *Z*_*i*_ = 1 otherwise).

Without making distributional assumption yet, we assume the pure cell type profile satisfies: *E*[*X*_*ik*_] = *μ*_*k*_ + ***Z*_*i*_**^*T*^ ***β*_*k*_**. Here *μ*_*k*_ represents the baseline profile for cell type *K*, and ***β*_*k*_** are coefficients associated with the covariates. The challenge is that *X*_*ik*_ is not directly observed. Instead, we observe signals that are mixtures of *X*_*ik*_’s. The observed data, denoted by *Y*_*i*_, is weighted average of *X*_*ik*_’s. For sample *i*, given the proportions ***θ*_*i*_**, we have

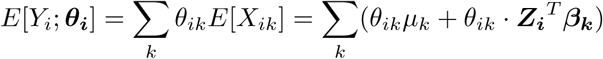

This is a typical linear model, with *μ*_*k*_ and ***β*_*k*_** as model parameters. The design includes mixing proportion as main effects, and mixing proportion by covariate interactions.

Assume we have ***Y*** from a total of *N* samples. Denote all observed data as ***Y*** = [*Y*_1_, *Y*_2_,…, *Y*_*N*_]^*T*^, the observed data can be described as a linear model: *E*[***Y***] = ***Wβ***, where

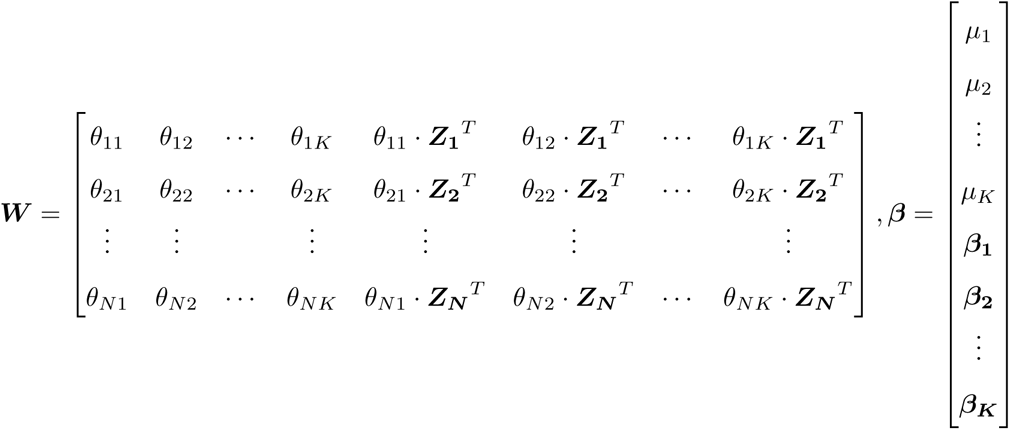

This setup captures the essence of several existing methods in an elegant classical linear model system. We will show in later sections that all existing methods are special cases of our model.

### Statistical inference for differential analysis

The above parameterization allows great flexibility in hypothesis testing for differential expression. Questions regarding a variety of expression changes can be answered by testing linear combinations of the regression coefficients. For example in a simple two-group (normal vs. disease) comparison setting, the covariate ***Z*_*i*_** reduces to an indicator, and each ***β*_*k*_** is a scalar. In this case, we have:

1. Testing the difference in cell type *k* between two conditions is *H*_0_: *β*_*k*_ = 0;
2. Testing the difference between cell types *p* and *q* in normal group is *H*_0_: *μ*_*p*_ *-μ*_*q*_ = 0.
3. Testing the difference between cell types *p* and *q* in disease group is *H*_0_: *μ*_*p*_ + *β*_*p*_ *-μ*_*q*_ *-β*_*q*_ = 0.
4. One can even test the higher order changes, for example, the difference of the changes between cell type *p* and cell type *q* in two conditions: *H*_0_: *β*_*p*_ - *β*_*q*_ = 0.

For multiple group comparison, for example, in eQTL studies where ***Z*_*i*_** has three levels (two degree of freedoms), F-test can be performed for cell type specific changes. We now can add distributional assumption on the observed data, for example, Gaussian for microarray or negative binomial for count data. The parameter estimation and statistical inference can be performed through linear model (for Gaussian data) or generalized linear model (for count data).

### Simulation setting

To evaluate the proposed method versus existing methods, and to examine the impact of different factors, we conduct a series of simulation studies. The simulated data are generated based on parameters estimated from real data, so that the simulation can well mimic the real data scenario. The general flow of simulation procedure is illustrated in Figure 2.

In the first step, we generate cell-type specific profiles (reference panel ***X***) based on the Immune Dataset. This dataset contains the gene expression profiles from four types of immune cells (Jurkat, IM-9, Raji, THP-1), each has measurements from three replicated samples [29]. For gene *g* in cell-type *k*, we first calculate the mean *μ*_*gk*_ and variance 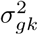 from the *log* expression values across the three replicated samples, where *g* = 1, *···, G*(*G* = 54657) and *k* = 1, *···, K*(*K* = 4). We assume each subject has a unique pure tissue profile ***X*_*i*_**, representing the biological variation among individuals even for pure cell type. ***X*_*i*_** is a matrix of *G* by *K*. For control samples, we simulate the *g*-th row and *k*-th column element of ***X*_*i*_** from a log-normal distribution with mean *μ*_*gk*_ and variance 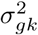 For cases, we first create some csDE genes. For each cell type, we randomly select 5% of the genes to be DE between case and control, half of them are up-regulated and half are down-regulated. The log fold changes for the DE genes are randomly drawn from *N* (1, 0.2^2^) for up-regulated genes and *N* (−1, 0.2^2^) for down-regulated genes. We then calculate the mean profiles for pure cell types in cases by adding the log fold change to *μ*_*gk*_. The variances 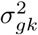 are kept unchanged for most simulations, except when we evaluate the impact of biological variance. In those simulations (results shown in Figure S2), we make the variances of pure tissue profiles in cases to be 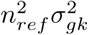, where we vary *n*_*ref*_ from 0.1 (small biological variance) to 2. The pure cell profiles ***X*_*i*_** for cases are then simulated from log-normal distribution. We simulate data for a total of *s*_1_ cases and *s*_2_ controls. Three selections of samples sizes (*s*_1_ = *s*_2_ = 50, 100, 500) are considered.

Next we simulate the mixing proportions ***θ*_*i*_**. For cases and controls we simulate from ***θ*_*i*_** ∼ *Dir*(***α*_1_**) and ***θ*_*i*_** ∼ *Dir*(***α*_0_**), respectively. The parameters ***α*_1_** and ***α*_0_** are based on a real dataset from Synapse.org (Synapse ID: syn6098424 [45]), which include 39 Alzheimer’s disease patients and 11 controls. We estimate the MLE of ***α*_1_** and ***α*_0_**, as 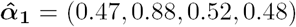 and 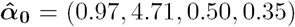 respectively. Using 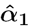 and 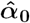, we generate *s*_1_ cases’ tissue proportions and *s*_2_ controls’ tissue proportions.

After reference panel ***X*_*i*_** and proportion ***θ*_*i*_** are obtained, the simulated measurements of subject *i* is ***Y*_*i*_** = ***X*_*i*_*θ*_*i*_** + ***E***. Both ***Y*_*i*_** and ***E*** are vectors of length *G*. Here ***E*** = {*ε*_*g*_, *g* = 1,…, *G*} represents the measurement error, and each element ε_*g*_ is simulated from N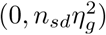. *n*_*sd*_ controls the level of technical noise. *n*_*sd*_ = 0.1, 1, 10 represent low, medium, and high noise levels. *η*_*g*_ is the standard deviation of measurement error for the *g*-th gene. To reflect the mean-variance dependence widely observed in expression data, we simulate *η*_*g*_ as a function of 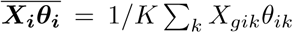. In this simulation, we use the relationship estimated from the Immune Dataset: 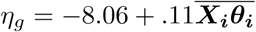.

When observed measurements are obtained, we use deconvolution methods to estimate the mixture proportion. Existing deconvolution methods, both reference-based (RB) algorithm, *lsfit* [29], and reference-free (RF) algorithm, *deconf* [37], are experimented with in this step. The proportion estimation uses the expression for a number of marker genes as input. For this purpose, we select 1000 genes with the largest variances of log expressions as potential marker genes. The expression of these genes can be directly fed into RF method *deconf* to estimate the proportions. For RB method *lsfit*, pure cell-type specific expressions for the marker genes are required as reference panel. We add measurement error to the true pure cell type profiles to generate the reference panel used in RB methods, to account for the fact that the reference in real data analysis are not known and have to be estimated from data. In real data applications, we use the same procedures to conduct RB and RF deconvolution as in simulation study.

### Model fitting in raw expression scale

Canonical methods for differential expression analysis have been based on log-transformed expression values, or have used log link function in generalized models on counts. There are a couple of reasons supporting this choice. First, from microarray intensity to sequencing counts, we have repeatedly observed the variance-mean dependence [46, 47, 48]. Genes with higher measurements on average, either in the form of microarray intensity or in sequencing counts, also appear to have greater variance. Second, measures of gene expression in tissue samples do not have units and only relative comparison in the form of ratio can be interpreted. Log transformation largely stabilizes the variance, making it relatively comparable across genes, and turns the ratio into differences. Stabilizing the variance, or making it comparable across genes, has another benefit in that a common prior on variance across genes can be used to obtain shrinkage estimates of gene-specific variance, by borrowing strength across genes in empirical Bayes approaches [46, 47, 48, 25]. Also, instead of testing a null hypothesis of ratio equating 1, testing the log ratio, i.e., the difference in log scale, becomes a familiar problem in linear models.

Since data mixing happens at the raw scale, model fitting is done in the same scale without taking the log transformation in our framework. Why does this still deliver good results? First, we note that in clinical studies with increasing sample sizes, we are no longer facing the problem of having so few replicates that the residual degree of freedom is too little to warrant a good estimate of variance. We demonstrate that in real data, given a gene, the distribution of raw expression across samples is approximately normal among replicates (Figure S7). Since the analysis is performed on the gene basis and no information is borrowed across genes, whether gene with higher expression also has higher variance is no longer relevant. Second, testing *β* = 0 in our model is equivalent of testing a ratio of 1, and we report 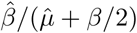 to keep the interpretation as a relative change.

### Several existing methods are special cases of the proposed method

A number of existing methods are available for detecting cell-type specific differential signals. Here we discuss the connections between three existing methods (cell-type specific differential methylation in human brain tissue [5], referred to as csDMHB; population-specific expression analysis (PSEA) [17]; and cell specific eQTL analysis [38], referred to as cseQTL) and the proposed method. We show that they are simplified version or special cases of our framework in testing cell-type specific differential signals. Note that none of these existing methods provide functionality for flexible hypothesis test, such as testing the difference among cell types in one condition as we showed in the Immune Dataset example.

#### Connection with csDMHB

csDMHB presented a statistical model that estimates differences in DNA methylation between two brain regions dorsolateral prefrontal cortex (D) and Hippocampal formation (H). Assume there are two cell types(NeuN+ and NeuN-) in the brain, they want to detect whether cell-type specific differences exist between the two brain regions. Following the notation from [5], *Y*_*i*_ is the observation for subject *i*. Indicator variable *X*_*i*_ = 1 if sample *i* from H and *X*_*i*_ = 0 otherwise. Their model is *E*(*Y*_*i*_) = *μ*_*D*,+_ + (*μ*_*D,-*_ *-μ*_*D*,+_)*π*_*i*_ + (*μ*_*H*,+_ *-μ*_*D*,+_)*X*_*i*_(1 - *π*_*i*_) + (*μ*_*H,-*_ - *μ*_*D,-*_)*X*_*i*_*π*_*i*_. To re-format this function into a linear model framework, they use *β*_0_, *β*_1_, *β*_2_, *β*_3_ to represent *μ*_*D*,+_, *μ*_*D,-*_ - *μ*_*D*,+_, *μ*_*H*,+_ - *μ*_*D*,+_, and *μ*_*H,-*_ - *μ*_*D,-*_ respectively.

Comparing csDMHB with our model, csDMHB is actually a simplified version of our framework with condition *K* = 2. Using our notation, when *K* = 2, we have *E*(*Y*_*i*_) = *θ*_*i*1_(*μ*_1_ + *β*_1_*Z*_*i*_) + *θ*_*i*2_(*μ*_2_ + *β*_2_*Z*_*i*_). Here *Z*_*i*_ is the indicator function that equals 1 if sample *i* from H and 0 otherwise. From this step, we can rearrange the term and build connections with csDMHB:

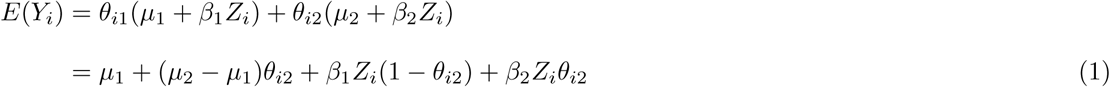

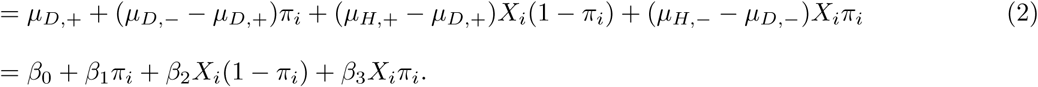

Transition from (1) to (2) implies *β*_0_, *β*_1_, *π*_*i*_, *β*_2_, *X*_*i*_, *β*_3_ in the notation of csDMHB corresponds to *μ*_1_, *μ*_2_-*μ*_1_, *θ*_*i*2_, *β*_1_, *Z*_*i*_, *β*_2_ in our notation system. They use the hypotheses *β*_2_ = 0 and *β*_3_ = 0 to test whether there is difference in NeuN+ methylation between D and H, and whether there is difference in NeuN-methylation between D and H respectively. This aligns with testing *β*_1_ = 0 and *β*_2_ = 0, which we proposes to test differences in cell type 1 and 2. Thus csDMHB is a simplified version of our method, for it only considers a mixture with two components. Moreover, it does not provide functionality to test the changes of different cell types under the same condition, e.g., difference between NeuN+ and NeuN-in D or H.

#### Connection with PSEA

PSEA presents a model formulation that detects the cell-type (or population) specific differential expressed genes with respect to their relative expression level calculated using cell-type specific marker gene expression level. To apply PSEA, one should first obtain a list of cell-type specific marker genes that express in cell-type *p*^*\**^ only. They assume 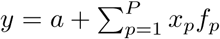 where a is background, *x*_*p*_ is the cell-type specific gene expression in cell type *p*, and *f*_*p*_ is the mixing proportion.

By using cell-type specific marker gene *x*_*p*_^*\**^, *f*_*p*_^*^ = (*y*_*p*_^*^ - *a*)*/x*_*p*_ or its approximation *f*_*p*_^*^ = *y*_*p*_^*^ /*x*_*p*_ is used as surrogate for mixture proportion. Thus the PSEA model becomes 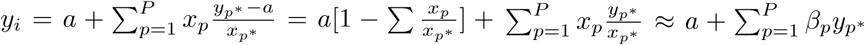. Here 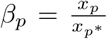. For two group comparison, PSEA uses an indicator variable *d*_*i*_ with *d*_*i*_ = 0 for samples in group 1 and *d*_*i*_ = 1 for samples in group 2. The model is represented as

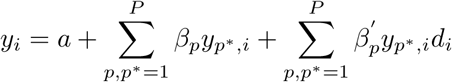

Comparing PSEA to our method, we find the ideas are very similar and our formulation is the generalized version of PSEA. Using our notation,

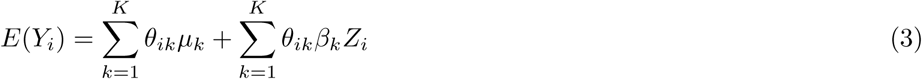

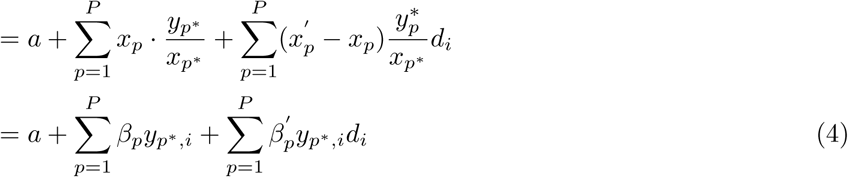

Transition from (3) to (4) suggests *β*_*p*_, 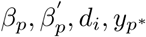, *d*_*i*_, *y*_*p*_^*^ correspond to *μ*_*k*_*/X*_*marker*_, *β*_*k*_*/X*_*marker*_, *Z*_*i*_, *θ*_*ik*_·*X*_*marker*_ respectively. Our method does not have marker gene expressionss in the formula since we assume the mixing proportions *θ*_*ik*_ are known, so we just use *X*_*marker*_ to symbolically represent the relative relationship. Note that the background term *a* is assumed omissible in PSEA and is absorbed into *μ*_1_ in our notation system. PSEA uses 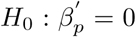 to test the null hypothesis of no difference between the *p*-th cell types in the two sample groups, which aligns with testing *H*_0_: *β*_*k*_ = 0 for detecting the difference between *k*-th cell types in our notation system.

Similarly in three-group comparison, PSEA has the model

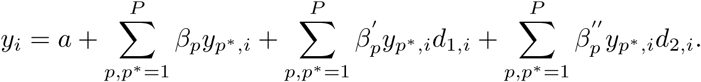

While in our notation system, we only need to specify covariate **Z_i_** as a vector with **Z_i_** = (0, 0)^*T*^ for subjects from group 1, (1, 0)^*T*^ for subjects from group 2, and (0, 1)^*T*^ for subject from group 3, then the correspondence relationship between PSEA and our model still holds. Thus, PSEA is similar to our method for detecting the expression changes for a specific cell type among different groups. It does not have functionality to test differential expression among different cell types in the same group, or higher order changes.

#### Connection with cseQTL

cseQTL expands the typical linear model to detect cell-type specific effects using expression quantitative trait loci (eQTL) datasets generated from whole tissue. Instead of using mixture proportions as the previous mentioned methods, cseQTL creates cell-type specific proxy for cell types of interest through correlation-based marker selection process. They treat the cell-type specific proxy as a covariate and add the main effect for the covariate and interaction term of covariate and genotype. Their model can be written as *Y* ≈ *I* + *β*_1_ × *G* + *β*_2_ × *P* + *β*_3_ × *P*: *G* + *e* where *Y* is the gene expression, *G* is the genotype information, *P* is the cell-type specific proxy, *P*: *G* is the interaction term between proxy and the genotype. The parameter of interest for testing is *β*_3_.

Comparing the cseQTL model with the existing methods and our proposed method, we find the cseQTL model is very similar to the formulation of csDMGHB and can be seen as a special case of method with *K* = 2. The fact that cseQTL only uses the proxy for one cell type in the formulation implies that they assume only two cell types (cell type of interest and cell type not of interest) in the analysis of each cell type. We can build a connection with cseQTL model as

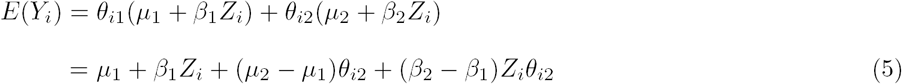

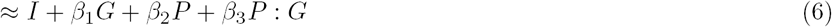

We use ‘≈’ to follow the notation utilized in the cseQTL paper [38] which indicates that cell-type proxy is used and the formulation just gives an approximation for single marker *cis-*eQTL mapping. The transition from (5) to (6) implies that we can build correspondence relationship between *β*_1_, *β*_2_, *β*_3_ in cseQTL’s notation and and *β*_1_, *μ*_2_ - *μ*_1_, *β*_2_ - *β*_1_ in our notation.

## Author’s contributions

HW and ZW proposed the statistical model. HW and ZL designed and performed the statistical simulation and real data analyses. The manuscript was written by ZL, ZW, and HW. All authors read and approved the final manuscript.

## Additional Files

Additional file 1: Supplementary.pdf

